# Decapitated body intelligence (DBI) in cricket *Gryllodes sigillatus*

**DOI:** 10.1101/2023.07.01.547349

**Authors:** Tongyao Xue, Hewei Yang, Wen Wu, Qinghao Wang

## Abstract

**Background:** intelligence is a highly complex problem. The decapitated insect exhibits various behaviors, indicating that they may have intelligence. Recent studies have reported organoid intelligence (OI) that lab-grown organoids also possess intelligence. This study investigated the response of decapitated crickets to different chemical stimuli to determine whether decapitated crickets have intelligence.

**Methods:** We used deionized water, NaCl, sucrose, and sodium hypochlorite to stimulate the front legs of the decapitated crickets and lesion the T1-T2 nerve connection of the thoracic ganglia. The behavioral response types of their forelegs were recorded. Reaction time, incidence rate, and total number of responses were calculated and analyzed.

**Results:** The decapitated crickets exhibit four types of responses: leg extension, leg withdrawal, leg lift, and jump. The reaction time and incidence rate varied depending on the type and concentration of the stimulant solution. The total number of responses gradually increased with deionized water, NaCl, sucrose, and sodium hypochlorite. The lesion experiments further revealed that the only T1 thoracic ganglion can control stimulating behavior.

**Conclusion:** The decapitated crickets possess the intelligence of perceiving stimuli and taking corresponding action. We called it decapitated body intelligence (DBI) or extrabrain intelligence (EI), which suggests that intelligence is not localized exclusively in the brain but also resides in insect ganglia. This finding opens up new ideas and avenues for the study of intelligence and the brain.

## 1. introduction

Intelligence has been explored in a wide variety of research areas, such as cognitive psychology, neuroscience, and medicine^1–5^. However, our current understanding of intelligence is limited, and new methods are needed to further explore this subject. Numerous studies have demonstrated that insects that have been decapitated can still respond to external stimuli and display various behaviors. Decapitated Drosophila have been observed to exhibit various behaviors such as movement, walking, flight, grooming, and sexually dimorphic behaviors in response to light, chemicals, biogenic amines, and stimulant drugs^6–9^. Shai Israel et al^10^ verified that decapitated flies can walk backward. Decapitated Drosophila can spontaneously exhibit some autonomous behaviors: combing, grasping, righting, and trembling^11^. The headless cockroaches showed escape turns, learning, and memory and reacted differently to cycloheximide and actinomycin D ^12–16^. In locusts, locust legs contract when stimulated by chemicals ^17^. Rogers and Newland reported that locust legs have chemoreceptors and can detect and respond to chemical stimuli ^18^. The decapitated cricket has also been studied in this area. The headless crickets also displayed a walk after defecation ^19^. A headless cricket will sing many of its songs^20^. The cricket’s front legs, which have sensory function, are most likely to be retracted by the stimulation of sodium chloride ^21^.

These studies indicate that some insects can survive for a period of time and respond to external stimuli after decapitation. The perception of stimuli and responses are some of the characteristics of intelligence, which suggests these decapitated insects could still have intelligence. There recently was a report on organoid intelligence (OI): cultivated organoids possess intelligence^22–24^, which prompted us to explore a new intelligent research method using decapitated crickets: decapitated body intelligence (DBI).

In this experiment, we used decapitated crickets as experimental models and stimulated their forelegs with the different chemicals. Because receiving stimuli and responding accordingly is an intellectual characteristic of animals, we observed and recorded the behavioral response types of their forelegs to clarify whether decapitated crickets can perceive stimuli from different solutions and make autonomous responses, and determine whether they possess intelligence.

## 2. Materials and methods

### 2.1 Experimental Materials

#### 2.1.1 Test insects

adults of the white cricket (Gryllodes sigillatus), Shengda Cricket Company

#### 2.1.2 Test reagents

sodium hypochlorite (Fujian Weizhen Biotechnology Co., Ltd.); sucrose (Beifangzhengda Chemical Reagent); sodium chloride (Beifangzhengda Chemical Reagent); deionized water.

#### 2.1.3 Test equipment

zoom stereo microscope SZM7045B1, OPPO RENOZ mobile camera, Micro tweezers, glass needles, Micro scissors.

### 2.2 Experimental Method

#### 2.2.1 Grouping

sodium chloride (NaCl) group, sodium hypochlorite (SH) group, sucrose group, and deionized water (DW) group (control group).

#### 2.2.2 Solution preparation

The effective concentration ranges of each chemical were determined based on preliminary experiments and references^17,18^, Divided into three concentrations: high, medium, and low. All solutions were prepared using deionized water (DW). For NaCl, the concentrations used were 10 mmol/l, 50 mmol/l, and 100 mmol/l; for sucrose and sodium hypochlorite, the concentrations were 100 mmol/l, 250 mmol/l, and 500 mmol/l.

#### 2.2.3 Anesthesia and decapitation treatment

Crickets were given sufficient food and water before the operation to ensure their activity. The crickets in each group were taken out of the feeding cages, and their heads were submerged in ice water for 20 seconds for anesthetization before decapitation. After removal from cold anesthesia (30 minutes), the crickets were re-tested. The remaining crickets underwent the same procedure sequentially before experimentation.

#### 2.2.4 Solution stimulation treatment and recording

Each decapitated cricket was laid on a plate placed on the operating table, and the camera was used to record their response. The solution for each group was dripped on the left tarsus of the front legs of the decapitated cricket using a Micro syringe. The experimental video recorded the cricket’s response continuously for 1 minute and was saved. All solutions were administered using the same method.

#### 2.2.5 Lesions of Thoracic Connectives

Following the method of Pedro et al^25^ and Lin et al^26^ with slight modifications, the thoracic ganglia’s T1-T2 nerve connection was selectively lesioned using a BD needle under a microscope.

#### 2.2.6 Data processing

Load the 120 recorded videos into AI SHI PIN video editing software on a computer and use the slow play function to observe and calculate the cricket’s The type of response, reaction time, incidence rate and total number of responses after being stimulated by the solution within 10 s after administration. Record the data in an Excel spreadsheet and plot it accordingly.

## 3. Results

### 3.1. Types of response

Within 10 seconds after the solutions stimulated the left front legs of the decapitated crickets, the decapitated crickets displayed the following behaviors: leg withdrawal, leg extension, leg lift, and jump. (Fig.1.). The jump occurred only when the sodium hypochlorite concentration was high and the body was curled up and jumped away from the original position. The decapitated cricket also exhibited some rare behaviors: other leg movements, wingspan, and somersaults. The behavior of decapitated crickets exhibited flexibility and diversity.

**Fig. 1.**
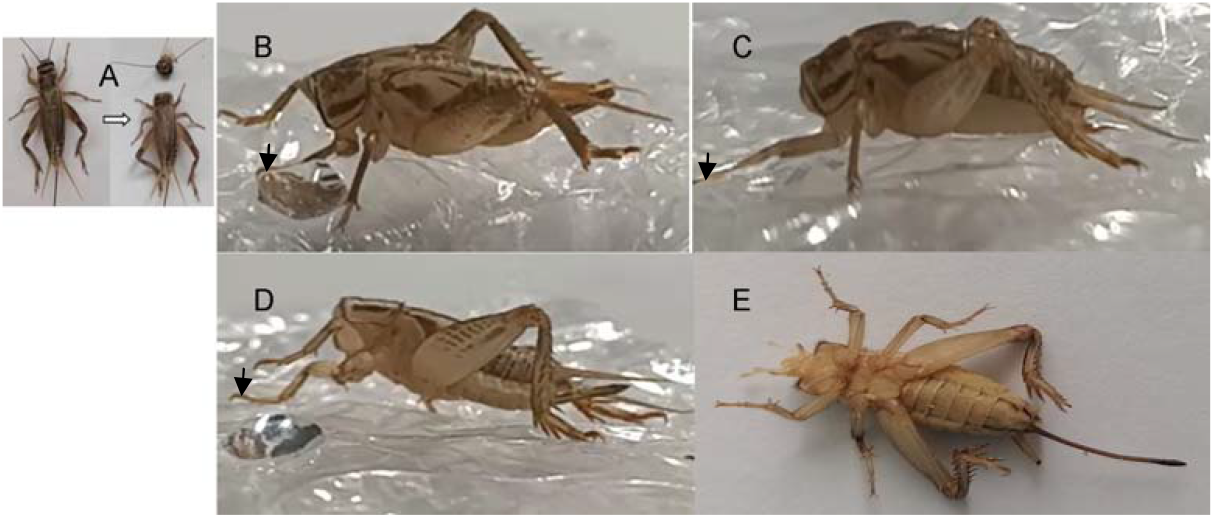
Behavioral types of the front leg of the decapitated cricket after the solutions stimulated. A. The cricket is decapitated B. The left front leg withdrawal C. The left front leg extension D. The left front leg lift E. The decapitated cricket jumps away, flipping its body with its belly facing upwards.

### 3.2. Reaction time

Reaction time is the time period from the first contact of a droplet with the tarsus to the start of the response. The average value of the reaction time of 12 crickets stimulated by each solution was recorded as the reaction time of this solution (Fig. 2). The response time of decapitated crickets varied depending on the type and concentration of the solution used. The response time decreased with increasing concentrations of SH, and the time required for the response was shortened when the concentration of SH was high. The reaction time was longest with a 100 mmol/l sucrose concentration, and 500 mmol/l sucrose had little gradient difference. However, the reaction time was longest with a 50 mmol/l NaCl concentration, and 10 mmol/l NaCl was the shortest.

**Fig. 2.**
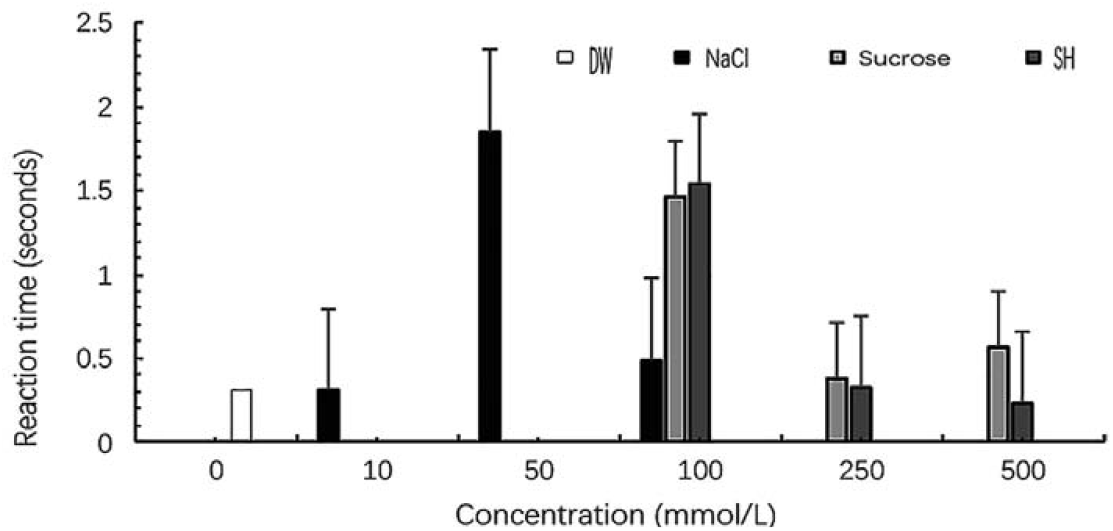
Reaction time of the front leg of the decapitated cricket. Mean±SD. Time is measured in units of seconds. Not Significant, Reaction time depend on both chemical identity and concentration.

**Fig. 3.**
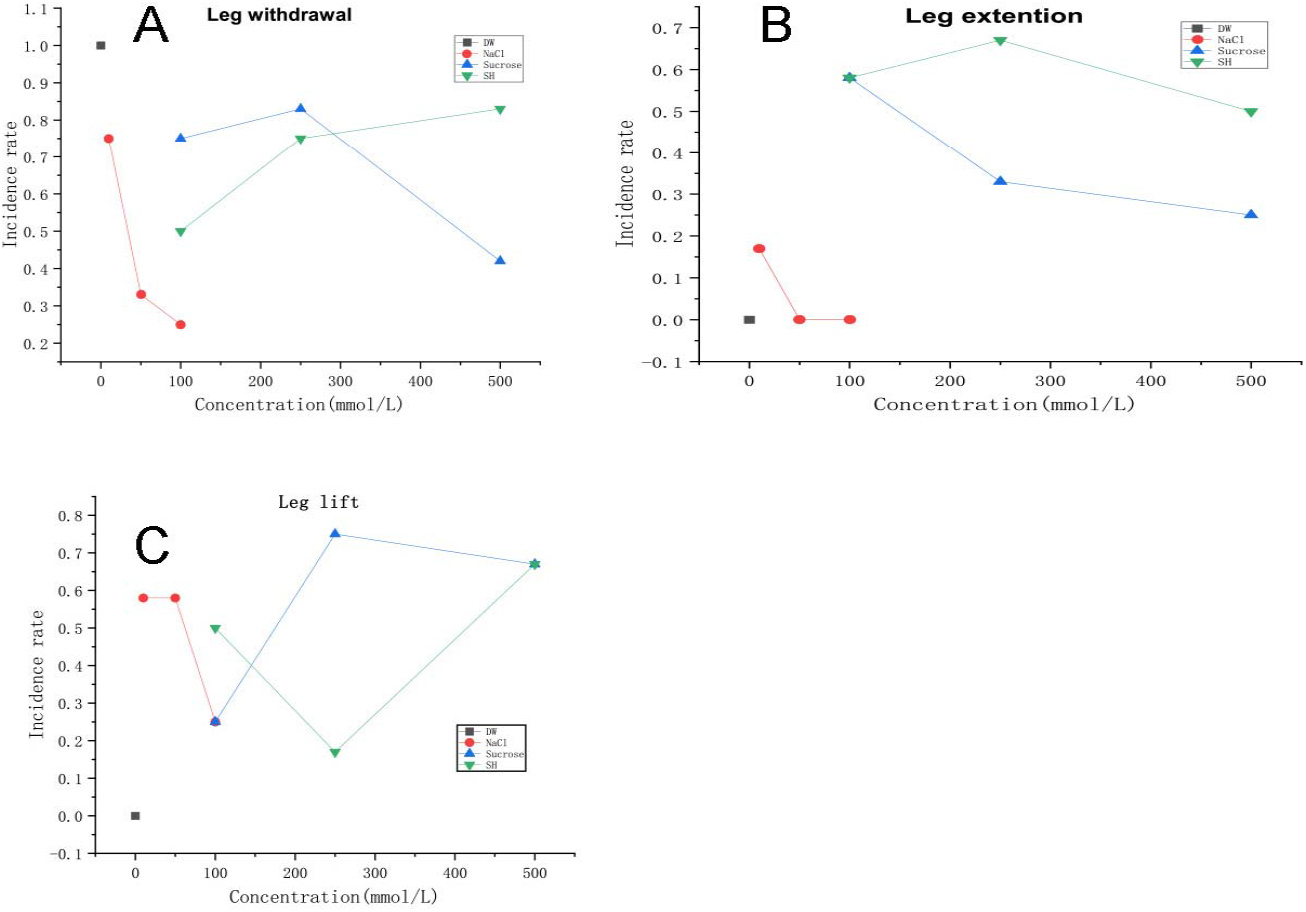
Incidence rate of the decapitated cricket. A. The incidence rate of leg extension; B, The incidence rate of leg retraction; C The incidence rate of leg lift. Incidence rate=response number/12(total), and time is within 10 seconds.

### 3.4. Total number of responses

The total number of responses refers to the sum of four behaviors (leg extension, leg contraction, leg lift, and jump) that occur within 10 seconds after the 12 decapitated crickets are stimulated by each solution. The total number of responses gradually increased with deionized water, NaCl, sucrose, and sodium hypochlorite (Fig. 4). In the total number of responses, sodium hypochlorite causes the most behavior 170 times, followed by sucrose 144 times, sodium chloride 63 times, and deionized water 12 times, showing that the number of behaviors is determined by chemical properties.

**Fig. 4.**
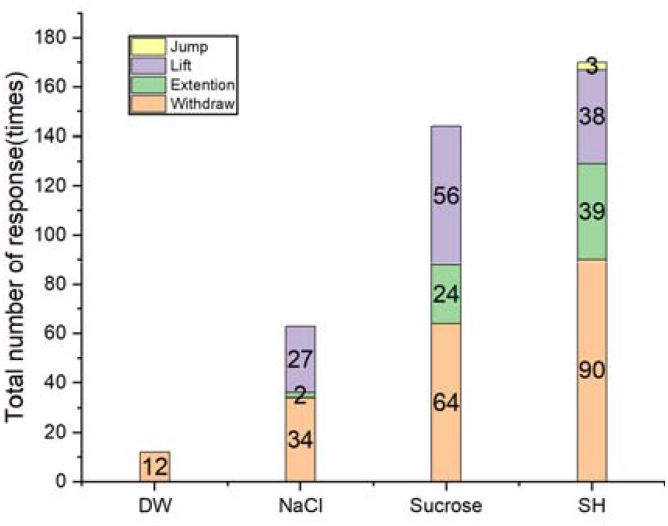
Total response number of each solution.

**Fig. 5.**
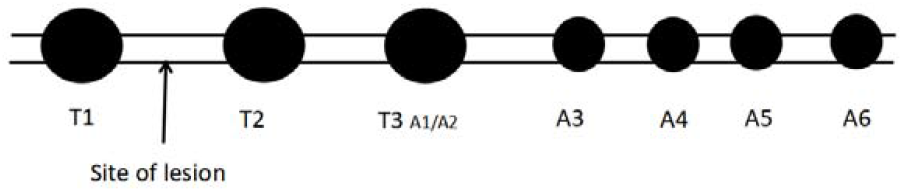
lesions of T1-T2 nerve connection. This image shows the ganglion of a cricket, and the nerve connection between T1 and T2 was lesioned.

### 3.5 Experiment of the lesion

A total of 12 decapitated crickets were used in this experiment. The T1-T2 nerve connection was lesioned, and the left front leg is stimulated by sodium chloride; the four reactions of extending, retracting, and lifting occur, but there is no jump. A single T1 ganglion has a dominant effect on chemical stimulation.

## 4. Discussion

This study conducted DBI experiments using decapitated crickets and examined the perception of decapitated crickets and their responses to different solution stimulations. The perception of stimuli and responses are some of the characteristics of intelligence. Our results show that intelligence is present in decapitated crickets and demonstrate that the intelligence is not localized exclusively in the brain.

Our results show that the decapitated crickets exhibited polymorphic behavior in response to different chemical stimuli, including leg retraction, extension, lifting, and jumping away. This result is analogous to that of Egeth and Yanagawa et al^6,7^. It is considered that because the stimulus is too strong, the severed cricket senses it and produces a violent response to quickly move away from it. The results also revealed that the thoracic ganglia can function independently of the brain, suggesting that intelligence perception and response can occur without the brain’s involvement.

The findings on reaction time and incidence rate reveal that the concentration and type of solution significantly impact the occurrence and frequency of special reactions. This indicates that these decapitated insects possess a level of intelligence that enables them to adapt to their environment and respond to different stimuli. Some studies have shown a correlation between reaction time and intelligence ^27,28^. This viewpoint also provides support for our research. The concentration of the solution does not always exhibit a linear relationship with the response; this phenomenon is analogous to that of Schubert ^29^. The total response number proves that headless crickets can distinguish different chemicals and make different levels of reactions.

The experiment with the lesion demonstrates that T1 alone responds to chemical stimulation, which indicates that this ganglion is responsible for the decapitated cricket’s autonomous decision-making ability. This also suggests that the thoracic ganglia can choose their own behaviors after receiving external stimuli without any input from the brain. A ganglion in an insect may be a mini-brain, which provides a new perspective for brain research.

The DBI in crickets offers researchers a unique opportunity to investigate the neural basis of intelligent behavior in a simplified model system. By using the individual ganglia of decapitated insects to solve difficult problems, researchers can avoid complex brain interactions. Importantly, the study provides a simple and convenient method to study intelligence mechanisms and create a simple model system of intelligence. These findings also provide practical implications for fields such as artificial intelligence and neuroscience and have several potential opportunities to explore fundamental questions about thought, decision-making, cognition, and consciousness.

The study has limitations. The stimulants used in the experiment were limited. Similarly, the study only investigated the response of the front legs, while the behavior of the other limbs remains unclear. Future research should consider expanding the range of stimulants, investigating other behavioral response types, and exploring other limbs’ contributions to DBI. Additionally, research should also explore more intellectual properties, essences, and mechanisms.

## Ethics approval and consent to participate

Not Applicable.

## Consent for publication

All authors approved the final manuscript and the submission to this journal.

## Availability of data and materials

The datasets generated and analyzed during the current study are available from the corresponding author on reasonable request.

## Authors’ contributions

Tongyao Xue: Investigation, Methodology, Data curation, Formal analysis, Validation, writing.

Hewei Yang: Investigation, Methodology, Data curation, writing-review.

Wen Wu: Investigation, Methodology.

Qinghao Wang: Conceptualization, Methodology, Validation, writing-review and editing, Supervision, Funding acquisition.

All authors read and approved the final manuscript.

## Acknowledgements

These studies were supported, in part, by the National Natural Science Foundation of China, 30672613.

